# microRNA165 and 166 modulate salt stress response of the Arabidopsis root

**DOI:** 10.1101/2022.11.25.517945

**Authors:** D. Scintu, E. Scacchi, F. Cazzaniga, F. Vinciarelli, M. De Vivo, R. Shtin, N. Svolacchia, G. Bertolotti, S. Unterholzener, M. Del Bianco, M. Timmermans, R. Di Mambro, S. Sabatini, P. Costantino, R. Dello Ioio

## Abstract

In plants, developmental plasticity allows for the modulation of organ growth in response to environmental cues. Being in contact with soil, roots are the first organ responding to soil abiotic stresses such as high salt concentration. In the root, plasticity relies on changes in the activity of the apical meristem, the region at the tip of the root where a set of self-renewing undifferentiated stem cells sustains growth. We show that salt stress promotes root meristem cells differentiation via reducing the dosage of the microRNAs miR165 and 166. By means of genetic, molecular and computational analysis, we show that the levels of miR165 and 166 respond to high salt concentration, and that miR165 and 166-dependent PHB modulation is fundamental for the response of root growth to this stress. Salt dependent reductions of miR165 and 166 causes rapid increase of the Arabidopsis homeobox protein PHABULOSA (PHB) expression and production of the root meristem pro-differentiation hormone cytokinin. Our data provide direct evidence of how the miRNA-dependent modulation of transcription factors dosage mediates plastic development in plants.

In plants, development must be both robust – to ensure appropriate growth - and plastic – to enable the adaptation to external cues. Plastic development largely depends on the modulation of gene expression, controlling the concentration of developmental factors, such as hormones, transcription factors (TFs) and signalling molecules (Garcia-Molina *et al*, 2013; Hofhuis & Heidstra, 2018; López-Ruiz *et al*, 2020; Schröder *et al*, 2021). A classic example of plant developmental plasticity is the adaptation of plant growth to high salt conditions, a stress that inhibits shoot and root development (Flowers *et al*, 1997). Roots are the first organs sensing salt concentration in soil, where high salt reduces meristem activity and root growth (Dinneny *et al*, 2008; Geng *et al*, 2013; Jiang *et al*, 2016). It has been suggested that the regulation of several plant hormones and miRNAs mediate the plant response to salt stress (Dolata *et al*, 2016; Geng *et al*, 2013; Iglesias *et al*, 2014; Jiang *et al*, 2016; Nishiyama *et al*, 2011; Yan *et al*, 2016). However, the molecular interplays mediating the adaptation of plant roots to salt stress are still vague. Post-embryonic root growth is supported by the activity of the root meristem, a region located at the root tip where self-renewing stem cells divide asymmetrically in the stem cell niche (SCN), originating transit-amplifying daughter cells that divide in the division zone (DZ) (Di Mambro *et al*, 2018). Once these cells reach a developmental boundary denominated transition zone (TZ), they stop dividing and start to elongate in the so-called elongation/differentiation zone (EDZ) (Di Mambro *et al*, 2018). A dynamic balance between cell division and cell differentiation ensures continuous root growth, maintaining a fixed number of cells in the DZ. Alterations in this dynamic equilibrium promote or inhibit root growth (Di Mambro *et al*, 2018; Salvi *et al*, 2020). microRNA molecules (miRNA) play a key role in the control of root meristem development (Bertolotti *et al*, 2021a; Skopelitis *et al*, 2012). Maturation of plant miRNAs depends on the activity of a multiprotein complex comprising the DICER-LIKE1 (DCL1), HYPONASTIC LEAVES1 (HYL1) and SERRATE (SE) proteins that cut pre-miRNA transcripts into 21 nucleotides mature miRNA (Yan *et al*, 2016). Among miRNAs, miR165 and 166 have been shown to be main regulator of root development (Carlsbecker *et al*, 2010; Dello Ioio *et al*, 2012). miR165 and miR166 are pleiotropic regulators of plant developmental processes. miR165 and 166 family consists of nine independent loci (*MIR165 A-B* and *MIR166 A-G*) that drive expression of pre-miR165 and 166 in different tissues and at different developmental stages (Miyashima *et al*, 2011). miR165/166 activity is crucial in the control of robust development, restricting the expression of the HOMEODOMAIN LEUCINE ZIPPER III (HD-ZIPIII), including PHABULOSA (PHB) and PHAVOLUTA (PHV), which are involved in root and shoot development, vascular growth, and leaf and embryo polarity (Carlsbecker *et al*, 2010; Dello Ioio *et al*, 2012; Di Ruocco *et al*, 2017; Grigg *et al*, 2009; McConnell *et al*, 2001; Skopelitis *et al*, 2017; Williams *et al*, 2005). In the root, miR165/166 regulate meristem homeostasis and radial patterning (Carlsbecker *et al*, 2010; Dello Ioio *et al*, 2012); pre-miR165a, pre-miR166a and b transcription is promoted by the SCARECROW (SCR) and SHORTROOT (SHR) transcription factors (Carlsbecker *et al*, 2010) and, thanks to the cell-to-cell mobility, mature miR165 and 166 forms diffuse to patterns both the root vasculature and the ground tissue (Carlsbecker *et al*, 2010; Miyashima *et al*, 2011; Skopelitis *et al*, 2018; Vatén *et al*, 2011; Bertolotti *et al*, 2021b). In the root meristem the miR165-166-PHB module promotes the synthesis of the plant hormone cytokinin, an important player in root developmental plasticity regulating cell differentiation rate of meristematic cells via the activation of the ARABIDOPSIS HISTIDINE KINASE3 (AHK3)/ARABIDOPSIS RESPONSE REGULATOR 1/12 (ARR1/12) pathway (Dello Ioio *et al*, 2007,2008). Here, we show that in response to salt stress miR165 and 166 modulates *PHB* expression to adjust root meristem activity. Salt exposure results in changes in cytokinin biosynthesis, which further regulates the miR165/166-PHB module. Hence, in addition to the above-described miRNA activity in controlling root robust development, we provide clear evidence that, in response to environmental cues, miRNAs are crucial also in the control of root plastic development, modulating the dosage of transcription factors.

## Results and Discussion

The growth of roots of Arabidopsis seedlings exposed to salt slows down after 5 hours of exposure to 150 mM NaCl (Geng *et al*, 2013) (SD1). We hypothesised that salt stress might inhibit root growth acting on meristem activity. To verify this, we analysed during time (5, 8, 16 and 24 hours) root meristem size in plants treated with 150 mM NaCl. A significant reduction in root meristem size was detected already after 5 hours of treatment (Fig 1 A-F). Analysis of stem cell and cell division markers such as *QC25* and *CYCB1;1:GUS* showed that salt exposure does not alter SCN and DZ activity (see SD1), suggesting that salt treatments mostly affect cell differentiation activity. To elucidate the molecular mechanisms behind root response to salt stress we first analysed the role of the miR165/166-PHB module.

**Fig. 1:**
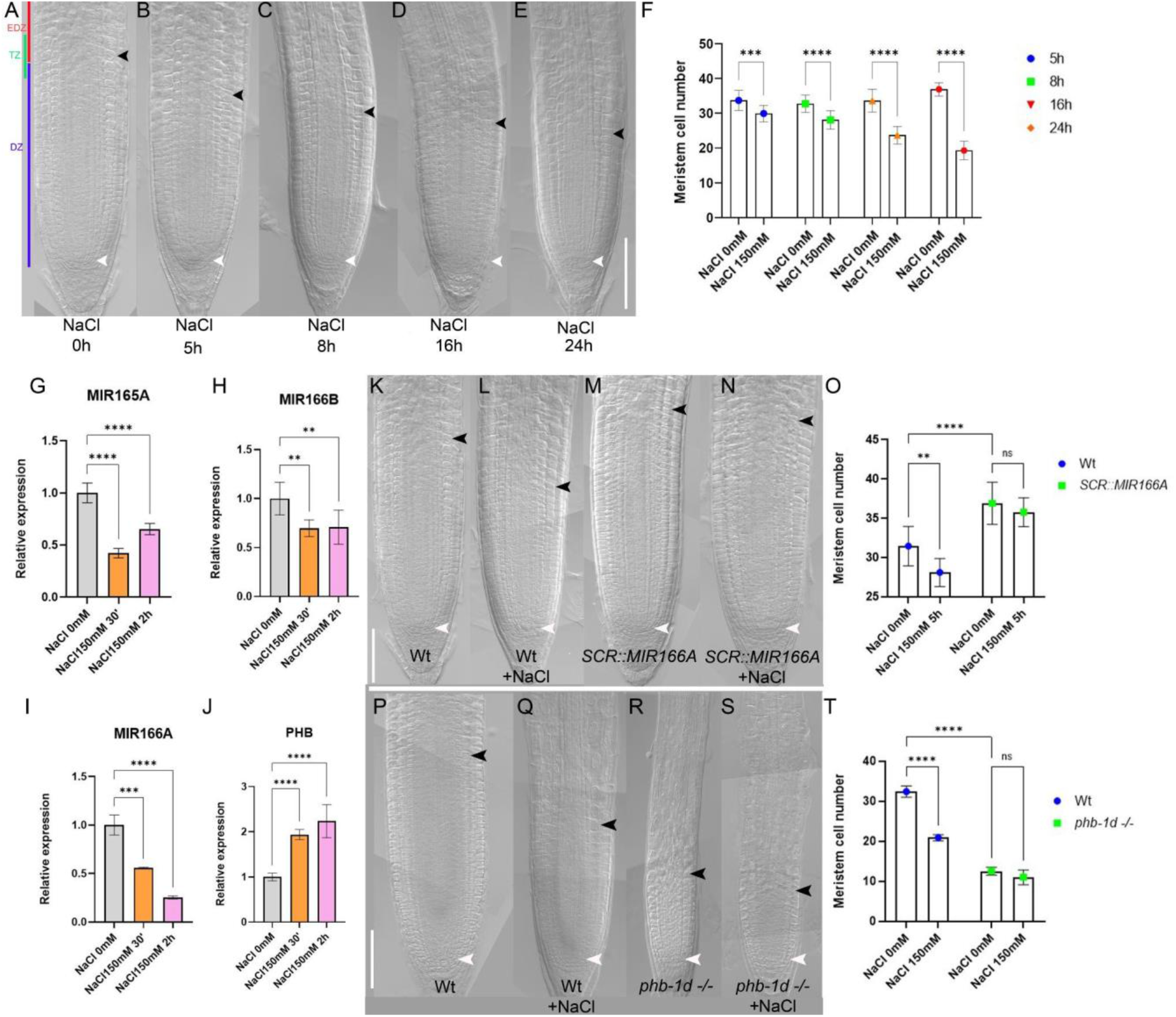
Salt stress inhibits root meristem activity regulating miR165 and 166 levels:= A-E) DIC of 5 days post germination (dpg) Wt root meristems exposed to 150 mM NaCl for 5, 8,16 and 24 hours (h). F) Root meristem cell number of Wt root meristems exposed to 150 mM NaCl for 5, 8,16 and 24 h. G-J) qRT–PCR analysis of preMIR165A (G), preMIR166A (H), preMIR166B (I) and PHB (J) RNA levels in the root tip of Wt plants upon 150 mM NaCl for 30 minutes and 2 h (**P<0.005, ***P < 0.001; **** P<0,0001 Student’s t-test; three technical replicates performed on three independent RNA batches). K-N) DIC of 5 dpg root meristems of Wt (K), Wt exposed to 150 mM NaCl for 5 h (L), *SCR::MIR166A* (M) and *SCR::MIR166A* exposed to 150 mM NaCl for 5 h (N). O) Root meristem cell number of Wt and *SCR::MIR166A* root meristems exposed to 150 mM NaCl for 5 h. N=30, Student’s t-test, Bar: SD. P-S) DIC of 5 dpg root meristems of Wt (P), Wt exposed to 150 mM NaCl for 5 h (Q) and *phb-1d -/-* (R) and *phb-1d -/-*(S). T) Root meristem cell number of Wt and *phb-1d-/-* root meristems exposed to 150 mM NaCl for 5 h. F,O, T) N=30, p < 0.00001; two-way ANOVA with Tukey’s multiple comparisons test, Bar: SD. A-E, K-N, P-S) Scale Bar 100μm, white arrowheads indicate the cortical stem cell, black arrowhead the TZ.

*MIR165A, MIR166A* and *B* are expressed in the Arabidopsis root meristem and control root meristem activity regulating *PHB* expression (Carlsbecker *et al*, 2010; Miyashima *et al*, 2011; Dello Ioio *et al*, 2012). We measured, via quantitative real time PCR (qRT), the mRNA level of PHB and pre-miRNA levels of miR165a, miR166a and miR166b after short-time exposures (2 hours) to 150 mM NaCl. Interestingly, roots subjected to salt treatment showed decreased levels of the pre-miRNA of miR165a, miR166a and 166b and higher transcription of PHB mRNA (Fig 1 G-J). Decrease in pre-miRNA occurs already after 30’ of salt exposure, preceding the increase of the PHB mRNA, which occurred only after two hours after salt treatments (Fig 1 G-J). This time lag suggests that NaCl specifically targets miR165a, miR166a and miR166b expression, which, in turn, regulates PHB mRNA levels.

To substantiate the primary role of miR165/166 in salt stress response, we investigated the response of plants expressing *MIR166A* under the control of the *SCR* promoter (*SCR::MIR166A*) to salt treatment. Since the *SCR* promoter is insensitive to salt stress and *SCR* is expressed in the endodermis as *MIR166 A, B* and *MIR65A* (Dinneny *et al*, 2008), SCR-driven *MIR166* expression should compensate the decrease of miR166 level caused by NaCl treatment (SD2). Indeed, plants carrying the *SCR::MIR166A* construct displayed longer root meristems than Wt and did not show any reduction in root meristem size even after 5 hours of 150 mM NaCL treatment (Fig 1 K-O), suggesting that the downregulation of miR165/166 is required for salt stress response. Concomitantly, the *phb phv* double loss-of-function mutant, that has a root meristem size resembling the one of *SCR::MIR166A* plants, showed no reduction in root meristem size after 5 hours of NaCl treatments (SD2).

Our data suggest that salt stress represses miR165 and 166 levels, causing a subsequent increase of the activity of HD-ZIPIII family members such as PHB. To corroborate this notion, we exposed *phb-1d* plants to 150 mM NaCl. In these plants, a mutation in the miR165 and 166 target site of the PHB gene prevents the miRNA-dependent PHB repression and causes increment in PHB expression responsible of generating a smaller meristem (McConnell *et al*, 2001). We reasoned that if salt stress goes specifically through miR165 and 166 regulation to adjust PHB levels, *phb-1d* plants should be resistant to salt exposure. As expected *phb-1d* plants were resistant to salt treatments, as their root meristem size did not vary upon NaCl exposure (Fig 1 P-T), These data suggest that NaCl specifically acts on miR165/6 decreasing their levels. Reduced miR165/6 levels cause an increase of PHB thus inducing root meristem arrest.

Thus, we assessed whether salt had a dose-dependent effect on miR165 and 166 levels and, consequently, on root meristem size. First, we analysed the response of the plant to different concentration of salt. We found that lower concentrations (100 mM NaCl) affect root meristem size less than 150 mM and 200 mM (Fig 2 A). To assess whether this dose-dependent developmental effect was indeed due to a dose-dependent modulation of miR165 and 166, we measured the levels of pre-miR165a, 166a, and 166b in root exposed for 2 hours to 100, 150 and 200 mM NaCl, via qRT-PCR. Indeed, we observed a strict correlation between the increase of salt concentrations and the levels of miR165 and 166 and the meristem size (Fig 2 B).

**Fig. 2:**
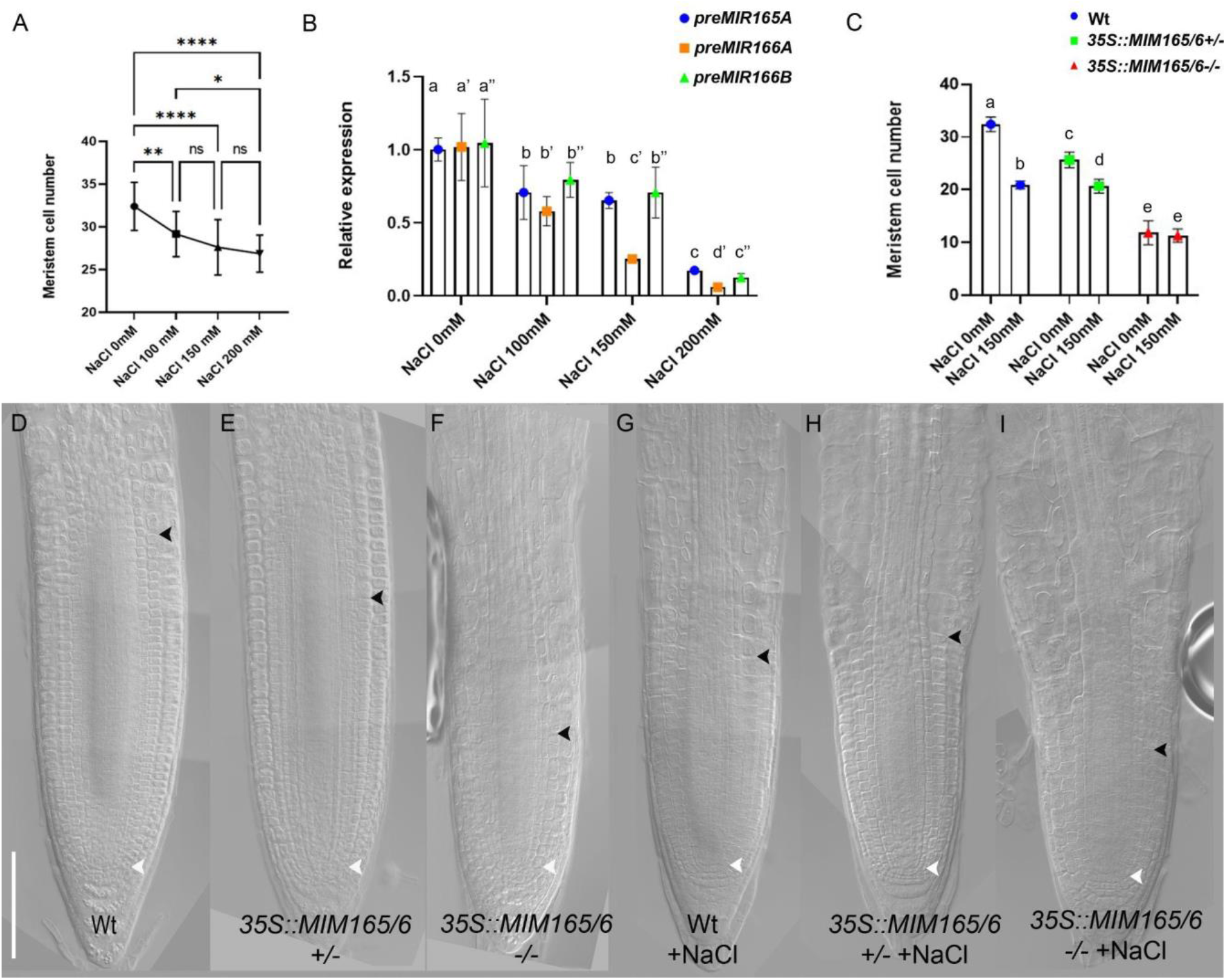
miR165 and 166 act in a dose dependent manner. A) Root meristem cell number of 5 dpg Wt root meristems exposed to 0, 100, 150 and 200 mM NaCl for 5 h. B) qRT–PCR analysis of *preMIR165A, preMIR166A, preMIR166B* RNA levels in the root tip of Wt plants upon 0, 100, 150 and 200 mM NaCl for5 h (three technical replicates performed on three independent RNA batches, p < 0.005; two-way ANOVA with Tukey’s multiple comparisons test. Different letters show statistical significance). C) Root meristem cell number of 5 dpg Wt, *35S::MIM165/6* +/-(het), *35S::MIM165/6* -/-(homo) root meristems exposed to 150 mM NaCl for 5 h. (N=30, two-way ANOVA with Tukey’s multiple comparisons test. Different letters show statistical significance). D-I) DIC of 5 dpg root meristems of Wt (D), *35::MIM165/6* +/-(E), *35S::MIM165/6* (F) and of Wt (G), *35::MIM165/6* +/-(H), *35S::MIM165/6* (I) root meristems exposed to 150 mM NaCl for 5 h. Scale Bar 100μm, white arrowheads indicate the cortical stem cell, black arrowhead the TZ.

To further corroborate the causal relation between miR165/166 levels and salt response of the root meristem, we manipulated miR165 and 166 levels exploiting the mimicry technology, which employs molecules that sequester and destroy miRNAs thus diminishing free miRNA (Todesco *et al*, 2010). We generated plants that constitutively express a mimicry targeting miR165 and miR166 (*35S::MIM165/166*). We analysed the root meristem size of *35S::MIM165/166* lines in both homozygosity (*35S::MIM165/166*) and heterozygosity (*35S::MIM165/166/-*) (Fig 2C-F). Both homozygote and heterozygote plants showed a shorter root meristem than Wt, but *35S::MIM165/166* homozygotes plants displayed a shorter root meristem than the *35S::MIM165/166/-* heterozygotes plants (Fig 2C-F). Moreover, *35S::MIM165/166* homozygote plants exposed to salt stress showed no root meristem size reduction, possibly because miR165/166 levels in these roots were already too low to allow for further repression (Fig 2 C-I). On the other hand, NaCl treatments reduced root meristem size in *35S::MIM165/166/-* heterozygote plants, although to a lesser extent than in Wt ones, presumably because of the salt-dependent downregulation of the residual miR165 and 166 levels (Fig 2 C-I). These data show that, upon salt stress, miR165 and 166 are responsible for the decrease in root meristem size in a salt-related dose-dependent manner.

Since miR165 and 166 modulates PHB levels, we set up to assess whether variations in PHB levels also affect root meristem size in a dose-dependent fashion. We first analysed the root phenotype of homozygous (*phb-1d*) and heterozygous (*phb-1d/+*) mutant plants. Both types of plants show shorter roots as compared to the Wt, with *phb-1d* roots being shorter than *phb-1d/+* ones (SD3).

The described *phb-1d* phenotype might in principle depend not only on a miR165 and 166 modulation, but also on a possible transcriptional regulation of PHB. Thus, we analysed the root meristem of plants where the ectopic transcription of a miRNA-insensitive version of PHB, fused to the GREEN FLUORESCENT PROTEIN (*phbmu-GFP*), was driven by a UAS/GAL4 transactivation system, bypassing a putative regulation dependent on the PHB endogenous promoter. Among the several available transactivation lines, the *Q0990* line was chosen because it drives expression only in the vascular tissue, where PHB is active. *Q0990>>phbmu-GFP* plants show a shorter root meristem than Wt. Notably, in analogy with *phb-1d* mutant analysis, *Q0990>>phbmu-GFP* homozygote lines displays a more severe phenotype than heterozygote ones (SD3). These data confirm the hypothesis that PHB-miR165/166 module is sufficient to cause variations in meristem size. We have previously showed that PHB regulates meristem size activating the transcription of the cytokinin biosynthesis *IPT7* gene (Dello Ioio *et al*, 2012). We therefore hypothesized that, in response to salt stress, PHB might modulate cytokinin levels, and hence root meristem size, via the regulation of *IPT7* expression. To assess this, we first exposed *ipt7* loss-of-function mutants to 5 hours of 150 mM NaCl and found that salt treatment does not affect the size of *ipt7* root meristems (Fig 3 A-E). This suggests that IPT7 is necessary to promote cytokinin-dependent cell differentiation in response to salt exposure. Then, we analysed the expression of *IPT7* in *phb-1d* and *phb1-d/+*, utilizing a nuclear-localized fluorescent transcriptional reporter of IPT7 (*pIPT7::NLS-3xGFP*)(Fig 3 F-I). We observed that IPT7 expression expands to the vasculature and endodermis in *phb-1d* and *phb1-d/+* compared to the Wt, where is mostly expressed in the columella and lateral root cap (Fig 3F-H). Moreover, a significantly higher GFP signal is detectable in the *phb-1d* homozygous line respect to the heterozygous one, thus suggesting that increased PHB levels are responsible for the enhanced IPT7 expression. Knowing that cytokinin via the AHK3/ARR1/12 pathway promotes cell differentiation inducing the expression of the IAA3/*SHORT HYPOCOTYL2* (*SHY2*) gene at the TZ (Dello Ioio *et al*, 2008), we assess a possible involvement of this circuit in salt stress response. We analysed the expression of *SHY2* in roots after 2 hours of NaCl exposure and found an induction of its expression (SD4). To assess whether salt-dependent regulation of *PHB* and *PHV* modulates *SHY2* expression at the TZ, we exposed *phb,phv,SHY2::GUS* plants to 150 mM NaCl. Interestingly, we were unable to detect any SHY2 expression at the TZ in this background, neither before nor after salt treatment (SD4), suggesting that the salt-dependent promotion of *SHY2* expression depends on PHB.

**Fig. 3:**
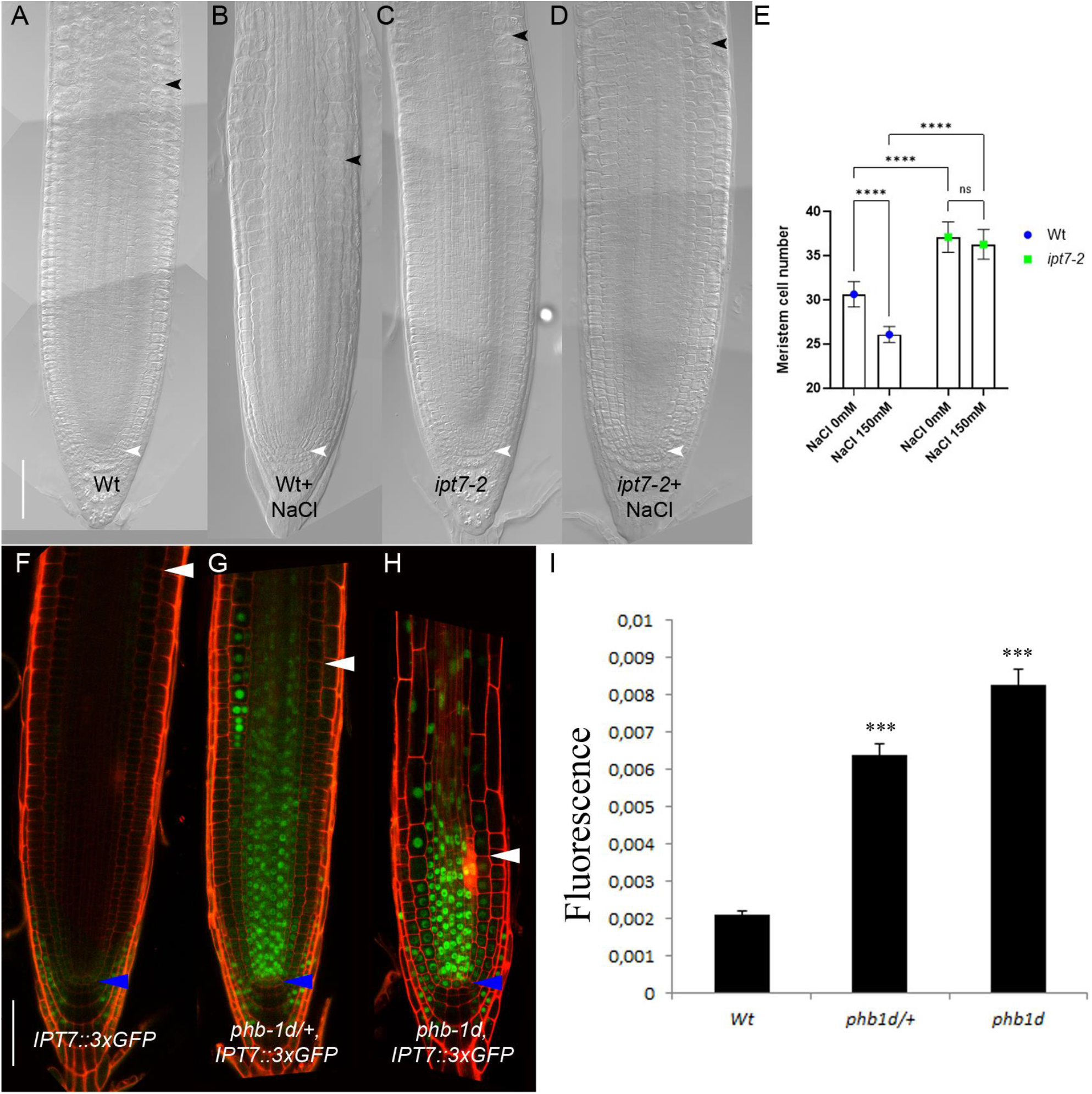
Salt stress inhibits root meristem activity via PHB/IPT7 module dosages. A-D) DIC of 5 dpg root meristems of Wt (A), Wt exposed to 5 h 150 mM NaCl treatments (B), *ipt7-2* (C), *ipt7-2* exposed to 150 mM NaCl treatments for 5 hours (D). Scale Bar 100 μm, white arrowheads indicate the cortical stem cell, black arrowhead the TZ. E) Root meristem cell number of 5 dpg Wt, *ipt7-2*, root meristems exposed to 150 mM NaCl for 5 h. (N=30, two-way ANOVA with Tukey’s multiple comparisons test. * shows statistical significance). F-H) Confocal images of 5 dpg root meristems of Wt (F), *phb-1d/+* (G) and *phb-1d* (H) carrying the construct *IPT7::3xGFP*. Scale Bar 50 μm, blue arrowheads indicate the cortical stem cell, white arrowhead the TZ. I) Quantification of *IPT7::3xGFP* fluorescence in the vascular of the root meristem of WT, *phb-1d +/-* and *phb-1d -/-*. N=5 two-way ANOVA with Tukey’s multiple comparisons test. * shows statistical significance.

These data suggest that salt exposure, by promoting cytokinin biosynthesis, enhances cytokinin-dependent cell differentiation activity at the TZ, hence, inhibition of root growth.

It was already reported that PHB is at the core of an incoherent loop, involving cytokinin and miR165/166: cytokinin directly inhibits PHB through ARR1, while activating it through the modulation of miR166 and miR165 to control root meristem development. This incoherent loop allows a rapid homeostatic regulation of PHB in response to fluctuations of cytokinin by maintaining an optimal threshold level of PHB necessary for proper root development (Dello Ioio *et al*, 2012). We thus questioned whether this circuit acting to keep the robust development of the root would maintain PHB homeostasis also in response to salt. To investigate how salt stress impacts on the behaviour of this circuit, we developed a mathematical model where the interactions between parameters are based *on in-vivo* experimental results. The model aims to understand how salt, cytokinin, miR165/6 and PHB, the components of the loop, react to changing salt concentrations, without determining the exact biochemical parameters of the system. According to the steady-state solutions of the model, as salt concentration increases miR165 and miR166 decreases, whereas PHB and cytokinin levels increase (Fig 4 A). For a more direct comparison with the experimental evidence, we simulated the system over time to determine the time course of the response of the circuit components to a salt-induced perturbation (Fig 4 B-C). According to the model, miR166/165 levels rapidly decrease after salt treatment. This causes the PHB level to rise, which results in cytokinin over-production, thus causing a decrease in meristem size and an inhibition of root growth. To understand how the levels of cytokinin are influenced by miR165/166-dependent PHB expression, we mimicked the miR165/166-insensitive PHB mutant (*phb-1d*) by performing two different time simulations of the model that, given the same initial conditions, differ for the inhibitory or non-inhibitory action of miR165/166 on PHB (dPHBmiR=0 in phb-d) (Fig 4 D). The model showed how the lack of PHB inhibition raises the levels of cytokinin, thus reflecting *phb-1d* smaller meristem phenotype. Moreover, the simulation of *phd-1d* mutant after salt exposure, validate the experimental observation that the mutant shows resistance to salt (Fig 4 E). In conclusion, our in-silico results confirm that the salt-mediated changes in mir166/165 levels are responsible for driving root developmental plasticity by regulating cytokinin production through PHB.

**Fig. 4:**
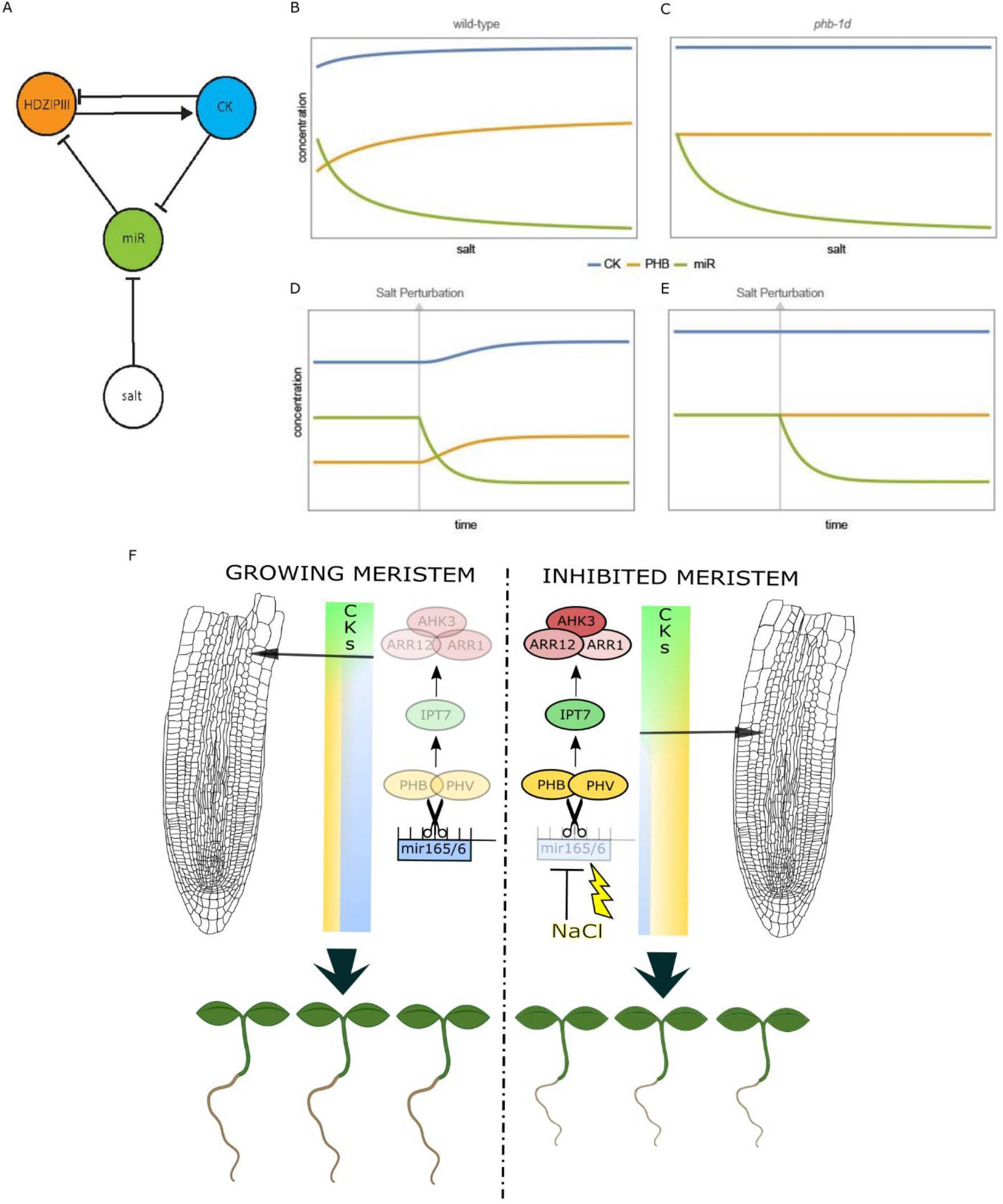
Salt-controlled miR165/166 levels are critical to determine root meristem size: A) Network topology of the model. B-E) Steady-State solutions of the model on a gradient of salt concentration in Wt B), and in *phb-d* D). Time-course simulating salt perturbation from a steady-state in wild type C), and in *phb-d* E). F) miR165/6 control root adaptation to salt stress: Cartoon depicting root meristem and root seedlings growing in standard conditions (left) and exposed to salt stress (right). In standard conditions levels of miR165 and 166 (cyan bar) maintain low PHB/PHV levels (yellow bar) and hence low cytokinin (CKs) activity via IPT7/AHK3/ARR1/12 circuit. In salt stress conditions (NaCl, yellow bolt) decreased levels of miR165/6 (cyan bar) cause increased PHB/PHV levels (yellow bar) promoting cytokinin (CKs) activity via IPT7/AHK3/ARR1/12 circuit. This process regulates the differentiation rate (diff) and hence whole root length. See text for details.

Altogether, our data show that miR165 and 166 control plastic development modulating PHB expression. This regulation is fundamental to adjust root plastic development in response to environmental cues such as salt exposure. In particular, salt stress decreases levels of miR165 and 166 resulting in an increase of PHB expression that induces a salt-dependent increase of cytokinin level via the induction of *IPT7* expression. Higher cytokinin levels at the TZ activates the AHK3/ARR1/12 pathway, which promotes the expression of *SHY2*, a negative regulator of the root putative morphogen auxin, triggering cell differentiation and represses root meristem activity in response to salt (Fig. 4 F)(Dello Ioio *et al*, 2008).

High cytokinin activity represses both the expression of PHB and miR165 and 166 (Dello Ioio *et al*, 2012). As previously suggested (Dello Ioio *et al*, 2012) and supported by our model, we posit that this regulation might help to maintain PHB levels in certain ranges in response to salt stress. Hence, this incoherent loop might help to provide a fast recovery of root growth after salt stress exposure, maintaining robust root development.

Recently, it has been shown that genes involved in auxin catabolism, such as *GRATCHEN HAGEN 3*.*17* (*GH3*.*17*), are induced by salt stress (Casanova-Sáez *et al*, 2022). *GH3*.*17* is a target of the AHK3/ARR1/12 module and, together with *SHY2*, generates an informative auxin minimum that triggers cell differentiation (Di Mambro *et al*, 2017, 2019). In future, it will be interesting to know whether salt dependent regulation of miR165 and 166 levels results in positioning the auxin minimum acting on *SHY2* and *GH3*.*17*.

Our results reveal that root adaptation to salt stress is driven by a modulation of miRNAs and miRNA target genes. Still to decipher is how and where salt is perceived in roots, and the details of the molecular pathways of salt stress response that lead to miR165 and 166 downregulation.

Salt-dependent regulation of miR165/166 might not only induce the regulation of meristem activity, but also coordinate other strategies of salt stress adaptation. Plants adapt to salt also generating xylem gaps in the root, stabilizing the DELLA gibberellin repressors (Augstein & Carlsbecker, 2022). Salt-dependent reduction of miR165/166 levels might coordinate this adaptation strategy by promoting PHB and its direct target GA2OX2, an enzyme involved in gibberellin catabolism (Bertolotti *et al*, 2021b), thus reducing gibberellin levels and stabilizing DELLA proteins. The utilization of halophyte models, such as *Eutrema salsugineum* (Cesarino *et al*, 2020), could help to clarify whether and how alterations in miR165/166/PHB module promotes tolerance to salt stress.

## Acknowledgments

We are grateful to Miltos Tsiantis for support through generation of materials and helpful discussions. To Paola Vittorioso, Davide Marzi and Patrizia Brunetti for thoughtful comments to the manuscript.

This work was supported by a Giovanni Armenise-Harvard Foundation mid-career development grant (to S.S.) and by Ministero dell’Istruzione, dell’Università e della Ricerca (MIUR).

## Conflict of Interest

The authors declare that the research was conducted in the absence of any commercial or financial relationships that could be construed as a potential conflict of interest.

## Materials and methods

### Plant Material and Stress Treatments

All mutants are in the Arabidopsis thaliana Columbia-0 (Col-0) ecotype background. For growth conditions, Arabidopsis seeds were surface sterilized, and seedlings were grown on one-half-strength Murashige and Skoog (MS) medium containing 0.8% agar at 22°C in long-day conditions (16-h-light/8-h-dark cycle) as previously described in (Perilli *et al*, 2010). Regarding stress treatments, 5 days old seedlings were grown on MS medium containing NaCl at different concentrations and for different periods of time as previously described in this paper.

*phb-13;phv-11* was previously described by (Carlsbecker *et al*, 2010), *35S::MIM165/6* by (Di Ruocco *et al*, 2017), phb1d and *phb,phv;SHY2::GUS* by (Dello Ioio *et al*, 2012). Enhancer trap line Q0990 was obtained from the NASC. *Q0990>>phbmu-GFP, phb1d;IPT7::3xGFP* were obtained by crossing. Homozygous and heterozygous lines were selected by phenotype.

### Generation and Characterization of Transgenic Plants

The IPT7::3xGFP plasmid was obtained as follow: 5.8 kb of IPT7 promoter driving nuclear 3xGFP were synthetized by GENEWIZ and inserted into PMLBART vectors via NotI flanking sites.

The *SCR::MIR166A* plasmid was obtained as follows: chpre-MIR166A was amplified from genomic DNA of Cardamine Oxford ecotype using specific primers: FW 5’-GGGGACAAGTTTGTACAAAAAAGCAGGCTGGGAGGAAGGAAGGGGCTTTCT-3’ REV 5’-GGGGACCACTTTGTACAAGAAAGCTGGGTGCCCTAATTAAATTGAGAAGAAGG-3’ and cloned in pDONR221 Gateway vector by BP recombination (Invitrogen). pDONRP4_P1-ChSCRp (Di Ruocco *et al*, 2017) and pDONR221-chpre-miR166A were recombined with pDONR P2R_P3-NOS into a pB7m34GW destination vector via LR reaction (Invitrogen; Karimi et al., 2002). ChSCRp has been utilized as it is expressed in the endodermis of *Arabidopsis thaliana* and it is not responsive to salt treatments (SD2A and B). Cardamine pre-miR166A (chpre-MIR166A) has been utilized to allow monitoring of endogenous Arabidopsis pre-miR166A response to salt as their sequences partially differ (SD2C, (Di Ruocco *et al*, 2017)). Cardamine miR166A and Arabidopsis miR166A mature forms are identical by sequence(Di Ruocco *et al*, 2017).

### Root length and meristem size analysis

Root meristem size for each plant was measured based on the number of cortex cells in a file extending from the quiescent center to the first elongated cortex cell excluded, as described previously(Perilli *et al*, 2010). The cortex is the most suitable tissue to count meristematic cells, as its single cell type composition shows a conserved number of cells among different roots. The boundary between dividing and differentiating cells for each tissue is called transition boundary (TB), while the region including the different transition boundaries is called transition zone (TZ; (Di Mambro *et al*, 2017)). For root MC analysis, root meristems of 5 days post germination (dpg) plants were analyzed utilizing a differential Interference Contrast (DIC) with Nomarski technology microscopy (Zeiss Axio Imager A2). Plants were mounted in a chloral hydrate solution (8:3:1 mixture of chloral hydrate:water:glycerol). Confocal images were obtained using a confocal laser scanning microscope (Zeiss LSM 780). For confocal laser scanning analysis, the cell wall was stained with 10 mM propidium iodide (Sigma-Aldrich). For each experiment, a minimum of 15 roots for three biological replicates were analyzed. Student’s t-test was used to determine statistical significance of these data (https://www.graphpad.com/quickcalcs/ttest1.cfm).

### RNA isolation and qRT-PCR

Total RNA was extracted from roots of 5 days old seedlings (both controls and treated with NaCl) using the NucleoSpin RNA Plus (Macherey-Nagel). Reverse-transcription was performed using using the SuperScript III First-Strand VILO cDNA Synthesis Kit (ThermoFisher Scientific). Quantitative RT-PCR (qRTPCR) analysis were performed using the gene-specific primers listed in Table 1

All the primers used were tested for their qPCR efficiency of 2-fold amplifications per cycle by qRT-PCR with the Standard curve method. PCR amplifications were carried out using the SensiFast SYBR Lo-Rox (Bioline) mix and they were monitored in real time with a 7500 Real Time PCR System (Applied Biosystems). UBIQUITINE 10 (UB10) amplification was used as housekeeper control and shown data are normalized to it. Data are expressed in 2-ΔΔct value.

Three technical replicates of qRT-PCR were performed on two independent RNA batches. Results were comparable in all the experiments and with the housekeeper. Student’s t test was performed to assess the significance of the differences between each sample and the control sample.

### GUS histochemical assay

β-Glucuronidase activity of transgenic lines carrying the GUS enzyme was assayed essentially as described in Moubayidin et al.12 using the β -glucoronidase substrate X-GlcA, (5-Bromo-4-chloro-3-indolyl-β-D-glucuronic acid, Duchefa) dissolved in DMSO. X-GlcA solution: 100 mM Na2HPO4, 100 mM NaH2PO4, 0.5 mM K3 K3 Fe(CN)6, 0.5 mM K4Fe(CN)6, 0.1 % Triton X-100 and 1 mg/ml X-GlcA. Seedlings were incubated at 37°C in the dark for an appropriate time allowing tissue staining depending on the GUS line assayed. Imaging was done using the Axio Imager A2 (Zeiss) microscopy. For each line and time-point, at least 15 roots were analyzed.

### Quantification and Statistical Analysis

Statistical analysis was performed using GraphPad (https://www.graphpad.com/). Error bars of all plots represent standard deviations (SD). The significance of the data was evaluated using the Student’s t test (*p < 0,05, p ** < 0,01, p*** < 0,001, NS Not Significant). All experiments have been performed in at least three replicas, using enough samples to ensure statistical significance.

### qRT PCR primers (Table 1)

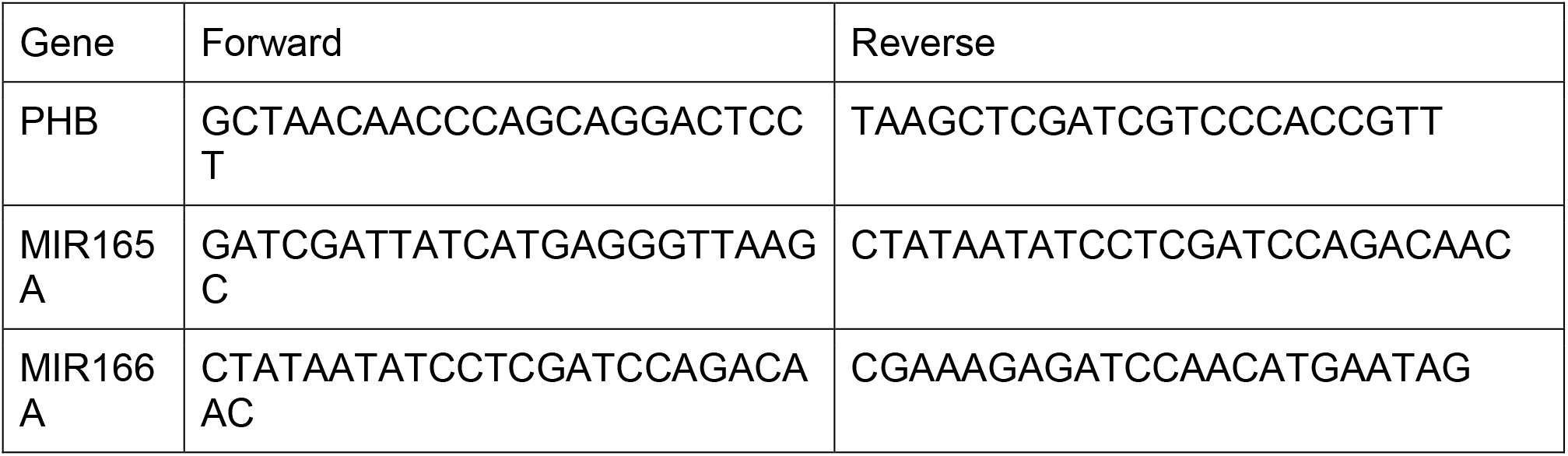
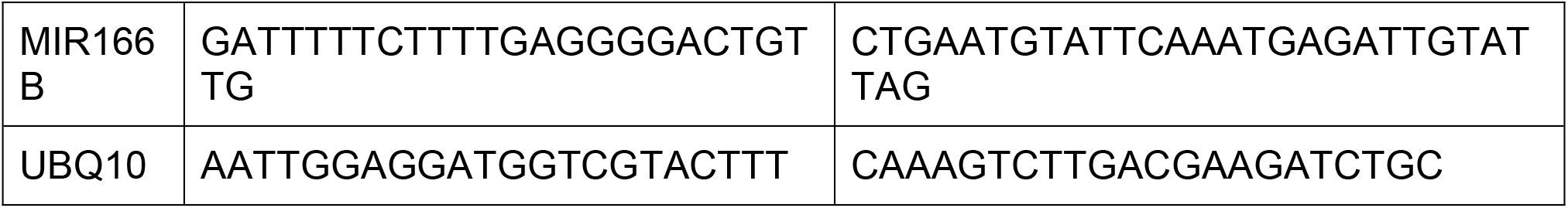

### Model Supplementary

To describe our data, we devised a simple mathematical model of the biochemistry of salt-mediated cytokinin regulation. The model, like the one described in (Dello Ioio *et al*, 2007), accounts for the time evolution of miR166, PHB and cytokinin relative to salt concentration. The model consists of the following set of coupled, first-order, ordinary differential equations:

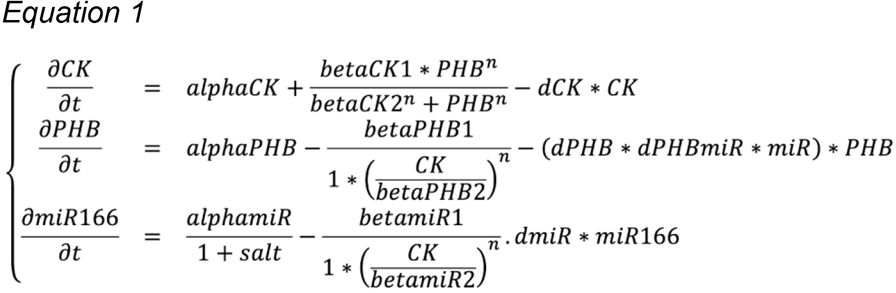

where CK, PHB and miR166 represent cytokinin, PHABULOSA and mir166/165 respectively. Adopting parameters as in (Dello Ioio *et al*, 2007) (Table), we used the software Wolfram Mathematica to solve the equations in (1) for the steady-state with the free parameter *salt* in a range between 0 to 10 (Figure). The system in (1) is solved numerically with parameters in Table using the Wolfram Mathematica NDSolve, which implements a Runge-Kutta method. To phenocopy the salt treatment of experiments on Arabidopsis plant, we use for, the value of the parameter salt equal to 0, and at time the parameter salt equal to 0.5 for a total *time* equal to 50 (Figure).

**Table.**

| Parameter | Details | Numerical Value |
| --- | --- | --- |
| alphaCK | Baseline of cytokinin production | 1 |
| betaCK1 | Maximum rate of PHB induced CK synthesis | 1 |
| betaCK2 | Apparent dissociation constant for PHB regulation of CK synthesis | 0.5 |
| dCK | Rate of CK degradation | 1 |
| alphaPHB | Baseline rate of PHB transcription | 1 |
| betaPHB1 | Maximum rate of CK-inhibited PHB transcription | 1 |
| betaPHB2 | Dissociation constant for CK regulation of PHB transcription | 0.4 |
| dPHB | Baseline rate of PHB degradation | 1 |
| dPHBmiR | Rate constant for mir165/6-induced PHB degradation | 1 (0 in phb-d mutant) |
| alphamiR | Baseline rate of mir165/6 transcription | 1 |
| betamiR1 | Maximum rate of CK-inhibited mir165/6 transcription | 1 |
| betamiR2 | Dissociation constant for CK regulation of mir165/6 transcription | 0.4 |
| dmiR | Rate of mir165/6 degradation |  |
| salt | Contribution of the salt on the rate of basal miR165/6 transcription | 0 or 0.5 |
| n | Hill coefficient | 1 |

## Supplemental Data

**SD1:**
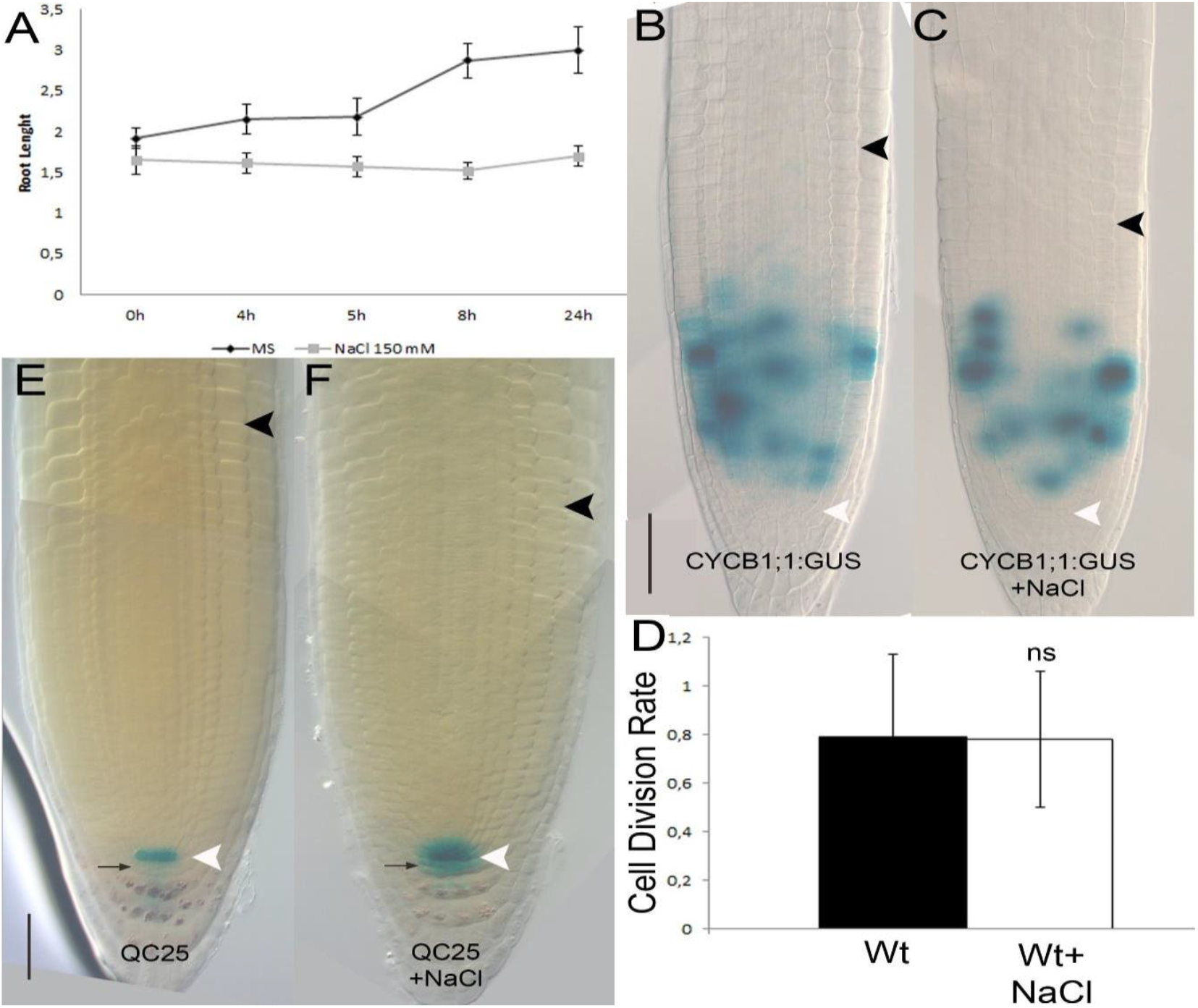
Salt stress inhibits root growth and meristem function regulating cell differentiation activity. (A) Root length over time of WT plants grown on standard MS medium in comparison to WT plants exposed to 150 mM NaCl. (Two Way ANOVA P<0,001, Error bars: SD; n =11) (B-C) DIC images of the histochemical b-glucuronidase (GUS) assay detecting CYCB1;1:GUS localization at 5dpg in a wild-type root meristem upon mock and NaCl (NaCl 150mM for 5 h) treatments. Scale Bar 100 μm, white arrowheads indicate the cortical stem cell, black arrowhead the TZ (n > 20). (D) Graph showing no reduction in cell division rate (gus stained cells/meristem cell number) after NaCl treatment (NaCl 150mM) in Wild-type plants (Col-0 background; ns not significant, Student’s t test; n > 20). (E-F) DIC images of the histochemical GUS assay performed in QC25::GUS plants upon mock and NaCl (NaCl 150mM for 5 h) treatments. Scale Bar 100 μm; white arrowheads indicate the cortical stem cell, black arrowhead the TZ.

**SD2:**
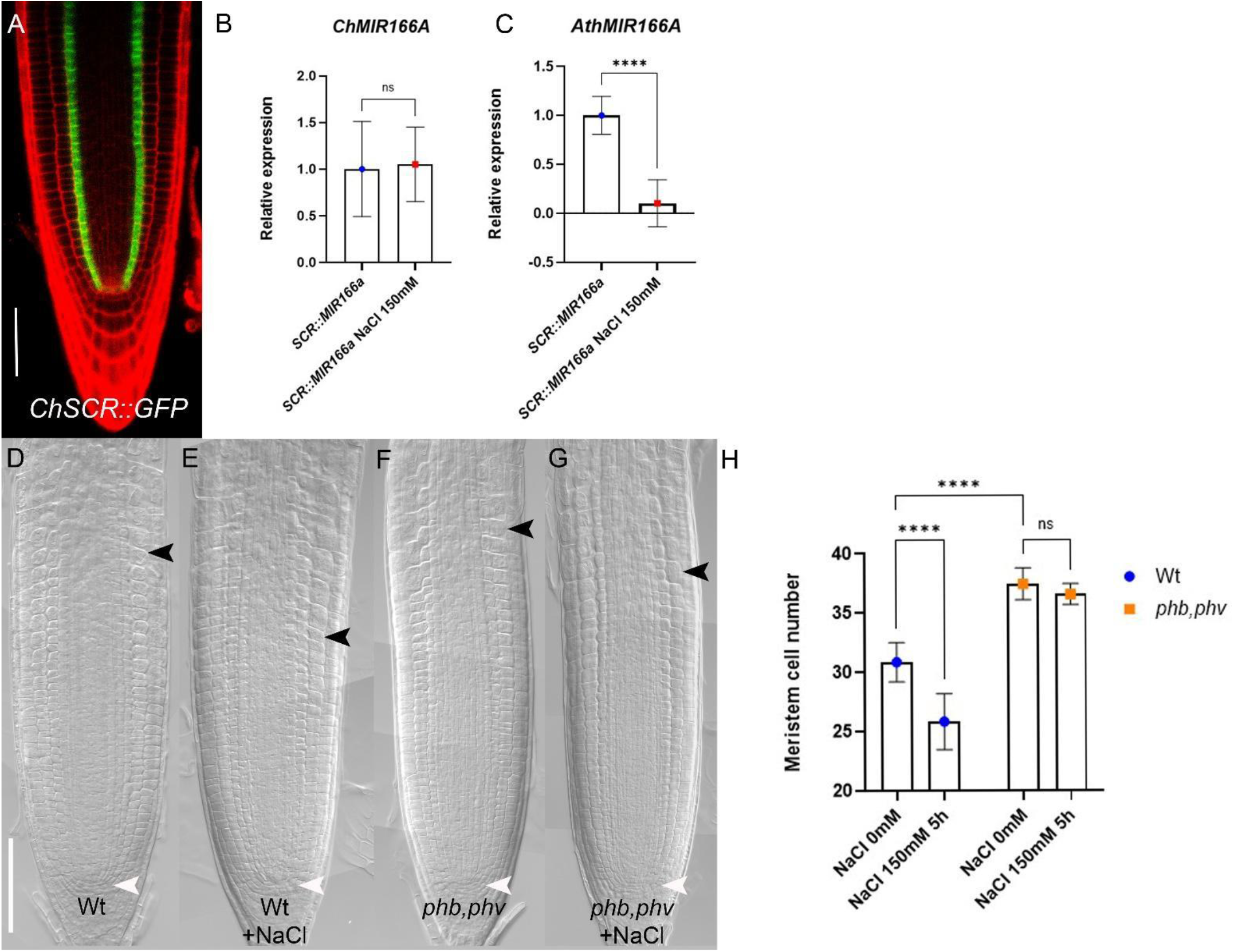
miR165 and 166 regulate root meristem size in response to salt stress (A) Expression of ChSCR::GFP in 5-DAG root meristem of Wt plants (Col-0 background). (B-C) qRT–PCR analysis of ChpreMIR166A (B) and AthpreMIR166A (C) RNA levels in the root tip of ChSCR::MIR166a plants upon 150 mM NaCl for 2 h (ns not significant ; ****p<0,0001, Student’s t test; three technical replicates performed on three independent RNA batches). (D-G) DIC images of 5 dpg root meristems of Wt (D), Wt exposed to 150 mM NaCl for 5 h (E), *phb,phv* (F) and *phb,phv* plants exposed to 150 mM NaCl for 5 h (G). Scale Bar 100 μm; white arrowheads indicate the cortical stem cell, black arrowhead the TZ (n>20). (H) Root meristem cell number of Wt and *phb,phv* root meristems exposed to 150 mM NaCl for 5 h (ns not significant, ****p<0.00001, two-way ANOVA with Tukey’s multiple comparisons test; Bar: SD; n>20).

**SD3:**
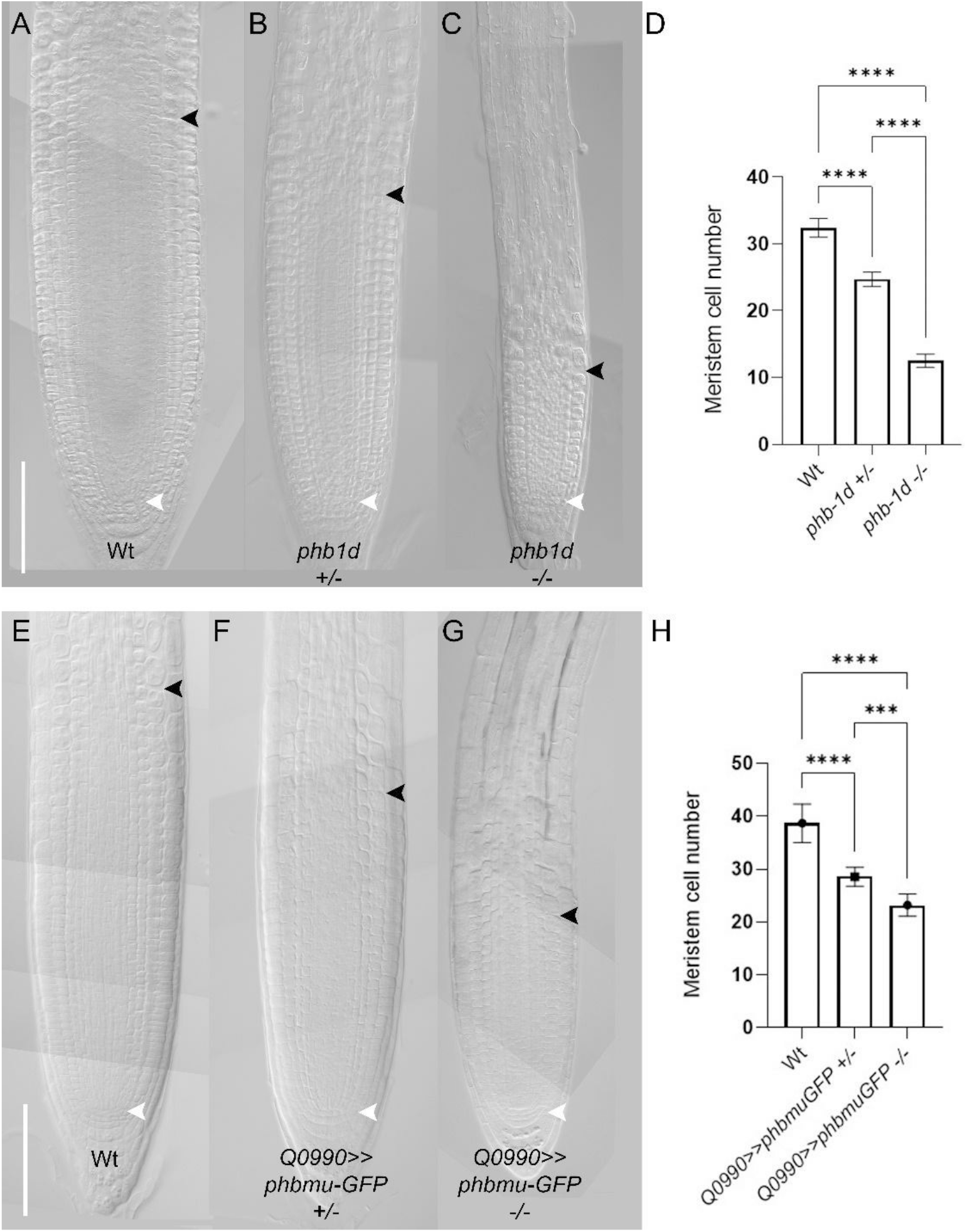
PHB controls root meristem size in a dose dependent manner. (A-C) DIC images of 5 dpg root meristems of Wt (A), phb1d heterozygous (B) and phb1d homozygous plants (C). Scale Bar 100 μm, white arrowheads indicate the cortical stem cell, black arrowhead the TZ (n>20). (D) Root meristem cell number of Wt, *phb1d* heterozygous and *phb1d* homozygous root meristems at 5 dpg (****p<0.00001, ***p<0,001, one-way ANOVA test; Bar: SD; n>20). (E-G)) DIC images of 5 dpg root meristems of Wt (E), *Q0990>>phbmu-GFP* heterozygous (F) and *Q0990>>phbmu-GFP* homozygous plants (G). Scale Bar 100 μm, white arrowheads indicate the cortical stem cell, black arrowhead the TZ (n>20). (H) Root meristem cell number of Wt, *Q0990>>phbmu-GFP* heterozygous and *Q0990>>phbmu-GFP* homozygous root meristems at 5 dpg (****p<0.00001, ***p<0,001, one-way ANOVA test; Bar: SD; n>20).

**SD4:**
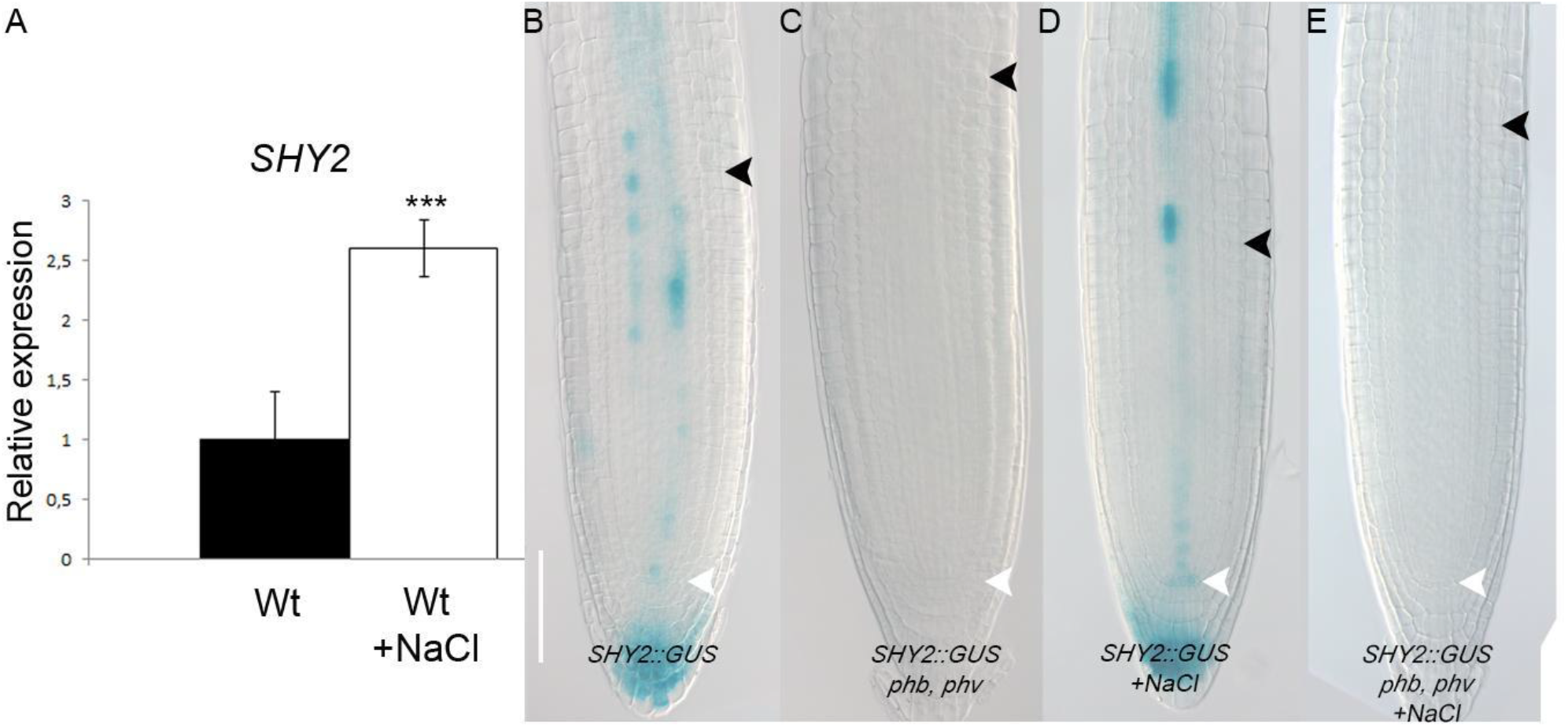
Salt stress promotes the expression of SHY2 in a PHB and PHV dependent manner. (A) qRT–PCR analysis of SHY2 RNA levels in the root tip of Wt plants upon 150 mM NaCl for 2 h (***p<0,001, Student’s t test; three technical replicates performed on three independent RNA batches). (B-E) DIC images of the histochemical b-glucuronidase (GUS) assay detecting SHY2::GUS expression at 5dpg in a wild-type root meristem upon mock(D) and NaCl (NaCl 150mM for 5 h; F) treatments. Scale Bar 100 μm, white arrowheads indicate the cortical stem cell, black arrowhead the TZ (n>20).

